# High resolution analysis of the cytosolic Ca^2+^ events in beta cell collectives in situ

**DOI:** 10.1101/2021.04.14.439796

**Authors:** Sandra Postić, Srdjan Sarikas, Johannes Pfabe, Viljem Pohorec, Lidija Križančić Bombek, Nastja Sluga, Maša Skelin Klemen, Jurij Dolenšek, Dean Korošak, Andraž Stožer, Carmella Evans-Molina, James D Johnson, Marjan Slak Rupnik

## Abstract

The release of peptide hormones is predominantly regulated by a transient increase in cytosolic Ca^2+^ concentration ([Ca^2+^]_c_). To trigger exocytosis, Ca^2+^ ions enter the cytosol from intracellular Ca^2+^ stores or from the extracellular space. The molecular events of late stages of exocytosis, and their dependence on [Ca^2+^]_c_, were extensively described in isolated single cells from various endocrine glands. Notably less work has been done on endocrine cells in situ to address the heterogeneity of [Ca^2+^]_c_ events contributing to a collective functional response of a gland. For this beta cell collectives in a pancreatic islet are particularly well suited as they are the smallest, experimentally manageable functional unit, where [Ca^2+^]_c_ dynamics can be simultaneously assessed on both cellular and collective level. Here we measured [Ca^2+^]_c_ transients across all relevant timescales, from a sub-second to a minute time range, using high-resolution imaging with low-affinity Ca^2+^ sensor. We quantified the recordings with a novel computational framework for semi-automatic image segmentation and [Ca^2+^]_c_ event identification. Our results demonstrate that under physiological conditions the duration of [Ca^2+^]_c_ events is variable, and segregated into 3 reproducible modes, sub-second, second and tens of seconds time range, and are a result of a progressive temporal summation of the shortest events. Using pharmacological tools we show that activation of intracellular Ca^2+^ receptors is both sufficient and necessary for glucose-dependent [Ca^2+^]_c_ oscillations in beta cell collectives, and that a subset of [Ca^2+^]_c_ events could be triggered even in the absence of Ca^2+^ influx across the plasma membrane. In aggregate, our experimental and analytical platform was able to readily address the involvement of intracellular Ca^2+^ receptors in shaping the heterogeneity of [Ca^2+^]_c_ responses in collectives of endocrine cells in situ.

## INTRODUCTION

Beta cell exposure to high glucose stimulates insulin release. Glucose entry into the beta cells through glucose transporters leads to a series of cellular events that trigger diffusion of Ca^2+^ ions into the cytosol. Resulting elevation in [Ca^2+^]_c_ ultimately produces a complex spatio-temporal pattern of [Ca^2+^]_c_ events, membrane potential changes, and exocytosis of insulin[1–3]. Single-cell recording of electrophysiological parameters are accepted as the gold standard approach to follow the plasma membrane events in a beta cell at a millisecond time scale[4, 5]. Electrical bursts composed of sub-second spikes were measured to have a plateau frequency of a few per minute[4] at high physiological glucose, while measured [Ca^2+^]_c_ oscillations were often orders of magnitude slower, with a period in a time scale of minutes[6, 7]. In the past, higher frequency and long-term recordings of [Ca^2+^]_c_ oscillations were limited due to relatively weak fluorescence signal of the classical fluorophores, their high affinity Ca^2+^ binding that yielded phase lags, slow sampling rates, and substantial bleaching[6–8].

There is growing evidence that intracellular Ca^2+^ release channels, namely the IP_3_ receptors (IP_3_R) and the ryanodine receptors (RYR), contribute to Ca^2+^-dependent insulin release in beta cells[9–12]. However, their exact role in physiological activation, activity and deactivation of beta cell is not clear[3]. A reactive oxygen species (ROS)- dependent mechanism was described to stimulate RYRs to support glucose-dependent insulin release in male rats[13]. Moreover, multiple studies suggest that disrupted activity of intracellular Ca^2+^ release channels may contribute to beta cell dysfunction and glucose intolerance [14–16].

To assess the contribution of intracellular Ca^2+^ release to [Ca^2+^]_c_ changes requires a much higher temporally resolved imaging. Fortunately, the advancement in synthesis of novel fluorescent Ca^2+^-sensing markers[17] solved most of the aforementioned technical issues enabling recording of [Ca^2+^]_c_ with millisecond temporal and high spatial resolution simultaneously in many cells with a possibility of repeated stimulation of the same cells over prolonged periods. High data volumes generated in this manner demanded development of an advanced data analysis, which is presented in this work. With these tools at hand, we conducted a series of experiments to revisit glucose-dependent beta cell activation, activity and deactivation, and with the help of pharmacological tools we reassessed the contribution of RYR intracellular Ca^2+^ release channels in these processes.

## RESULTS

### Imaging of beta cell [Ca^2+^]_c_ in pancreas slices at physiological glucose concentration

Pancreas tissue slices offer the opportunity to study beta cells in situ (Figure 1) and closer to physiological conditions compared to isolated islets and dispersed islet cells [18]. At sub-stimulatory glucose concentrations (6 mM), beta cells from adult mice displayed a stable resting [Ca^2+^]_c_, which can be seen during the pre-stimulatory and subsequent washout phases (Figs. 1 BD). From a baseline of 6 mM, a physiological stimulatory glucose concentration (8 mM) after a typical delay of solution exchange and glucose metabolism, triggered a biphasic [Ca^2+^]_c_ response (Figs. 1BD). This biphasic response is composed of so-called initial increase in [Ca^2+^]_c_, previously termed the transient or asynchronous phase, followed by a prolonged plateau phase (Figs. 1BD)[8, 19]. In both phases, we detected [Ca^2+^]_c_ events at different time scales, spanning from millisecond to a hundred of seconds range (Figs 1B,D-F,I-L). These [Ca^2+^]_c_ events show several levels of temporal summation following a self-similarity pattern, as long events were a temporal summation of series of short events, a feature that could be indicative of Ca^2+^-induced Ca^2+^ release-like (CICR) behaviour (Figure 1DH). In many aspects, the arrangement of short and long events reliably follows the electrical activity recorded in classical studies[4].

**Figure 1.**
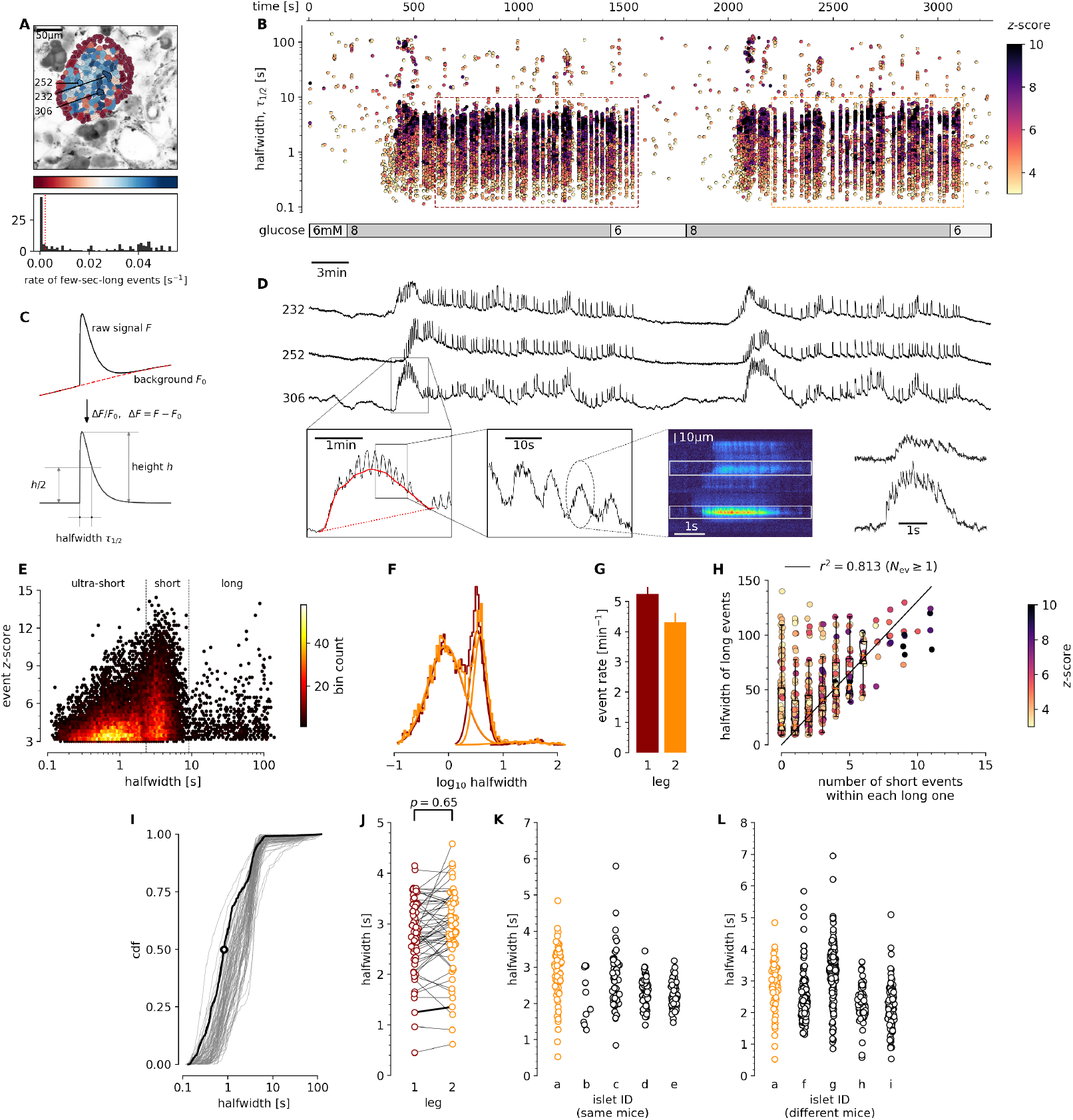
Quantification and analysis of [Ca^2+^]_c_ dynamics in physiological stimulatory glucose. **A**, Regions of interest (ROIs) obtained by our segmentation algorithm. The color indicates the number of events identified in the ROI trace, upon a high-pass filtering at 0.2Hz. We discarded ROIs with number of events below the threshold (red dashed line in the histogram in the lower panel). Indicated are the ROI numbers whose filtered traces correlate best with the average trace for the whole islet; the correlation coefficients being 0.835, 0.818 and 0.807 (from top down in D). **B**, Events’ halfwidth duration through time. Note the ranges of halfwidth duration occurring, the events’ synchronicity, and the phenotype’s reproducibility in sequential glucose stimulation. Color indicates the statistical significance in terms of *z*-score. Only events with a *z*-score higher than 3 were included. The stimulation protocol is indicated in the bar at the bottom of the pane. The plateau phases in both stimulations are indicated by a color coded rectangles. There is a prominent superposition of the short events on the plateau phases between the ROIs. **C**, An schematic of a transient event to describe the features of [Ca^2+^]_c_ events from background subtraction of the raw data, to height and halfwidth duration. Peak-point is the time of highest amplitude of an event. **D**, *(Upper Panel*) Time courses from ROIs indicated in A, exposed to a double stimulation protocol, and rebinned to 2Hz (recorded 20Hz). The abscissa is shared with B and is indicated there. *(Lower Panels*) Illustration of the compounding nature of transients. The left-most panel is a closeup of the trace above to highlight the long transient in red. In the second closeup we emphasize the structure within the long transient. The third panel shows data from a different experiment, in same conditions (8 mM glucose), recorded as a line scan. In the final panel, we plot the time course of the indicated spatial regions, to illustrate the structure of sub-second events. **E**, In a *z*-score vs. halfwidth density plot, the events clearly separated into three groups, which we named ultra-short, short and long, with a typical separation between the groups at 2 s and 8.5 s. The dominant time scale of the events had a halfwidth duration of 2-5 s. **F**, Normalized Gaussian fit through the logarithmic distribution of halfwidth duration for events for both first and second stimulation as indicated in B. **G**, The rate of events for the dominant short [Ca^2+^]_c_ time scale for both stimulations indicated. **H**, Evidence that the long events resulted from a progressive temporal summation of the (ultra-)short events. Some of the long events are less likely to contain substructure of short events, have lower *z*-scores, and contribute only little to the least square fit. **I**, Cumulative distribution frequency (cdf) of the halfwidth duration of dominant short events during the plateau phase of the first stimulation for a randomly selected ROI. Thick black line indicates the median distribution. **J**, Comparison between the halfwidth durations of events in individual ROIs during first and second glucose stimulation. **K**, Comparison of halfwidth durations of events from all ROIs from different islets of the same mouse. **L**, Comparison of halfwidth durations of events of all ROIs from islets of different mice.

The initial transient phase is characterized by sizeable delays between the activation of individual beta cell groups within an islet[20]. In the initial phase we recorded one or more large transient [Ca^2+^]_c_ events with a mean duration of tens of seconds (Figure 1D), and with or without discernible superimposed shorter [Ca^2+^]_c_ events. The long transient events were also occasionally recorded outside the initial phase. In less than 30 % of recorded islets or in a small fraction of the cells in a typical islet, long events were more prominent and were apparent in both initial as well as in plateau phase.

Meanwhile, during the plateau phase we recorded mostly [Ca^2+^]_c_ events with a mean halfwidth duration of a few seconds (Figure 1D-G), occasionally exhibiting temporal summation into slowly rising and decaying long events (Figs. 1DH). These short events were a dominant time scale in the control conditions at 8 mM glucose in all experiments. The short [Ca^2+^]_c_ events within the plateau phase were commonly well synchronised among the beta cells in an islet (Figs. 1BD). The short [Ca^2+^]_c_ events on the plateau phase were regenerative and could be stably recorded for hours at physiological glucose concentrations[21]. Due to the regenerative nature of the short [Ca^2+^]_c_ events, we could design a multiple stimulation protocol on the same slice. The protocols were designed in a way that in the initial section we tested the responsiveness to the stimulation with 8 mM glucose, in the second section we tested the effect of a specific pharmacological treatment on stimulation with 8 mM glucose, and in the last section we applied another 8 mM glucose control stimulation (Figure 3). A detailed inspection of the short [Ca^2+^]_c_ events in the plateau phase showed that they were compound events composed of even shorter [Ca^2+^]_c_ events with a mean halfwidth duration below one second (Figs. 1E, 2C, 5C). Similar ultra-short events were until now observed only as spikes on top of the bursts of the electrical activity, and were attributed to Ca^2+^ action potentials[22–24], mediated by the dominant contribution of L-type VACCs[25]. Our approach offered insight into [Ca^2+^]_c_ events on times scales spanning several orders of magnitude (Figure 1E), with temporal resolution comparable to that achieved with electrophysiological approach, and with an upgrade of simultaneously visualising the activity of all islet cells in an optical plane permitting the interrogation of a collective behaviour in situ (Figure 1A).

**Figure 2.**
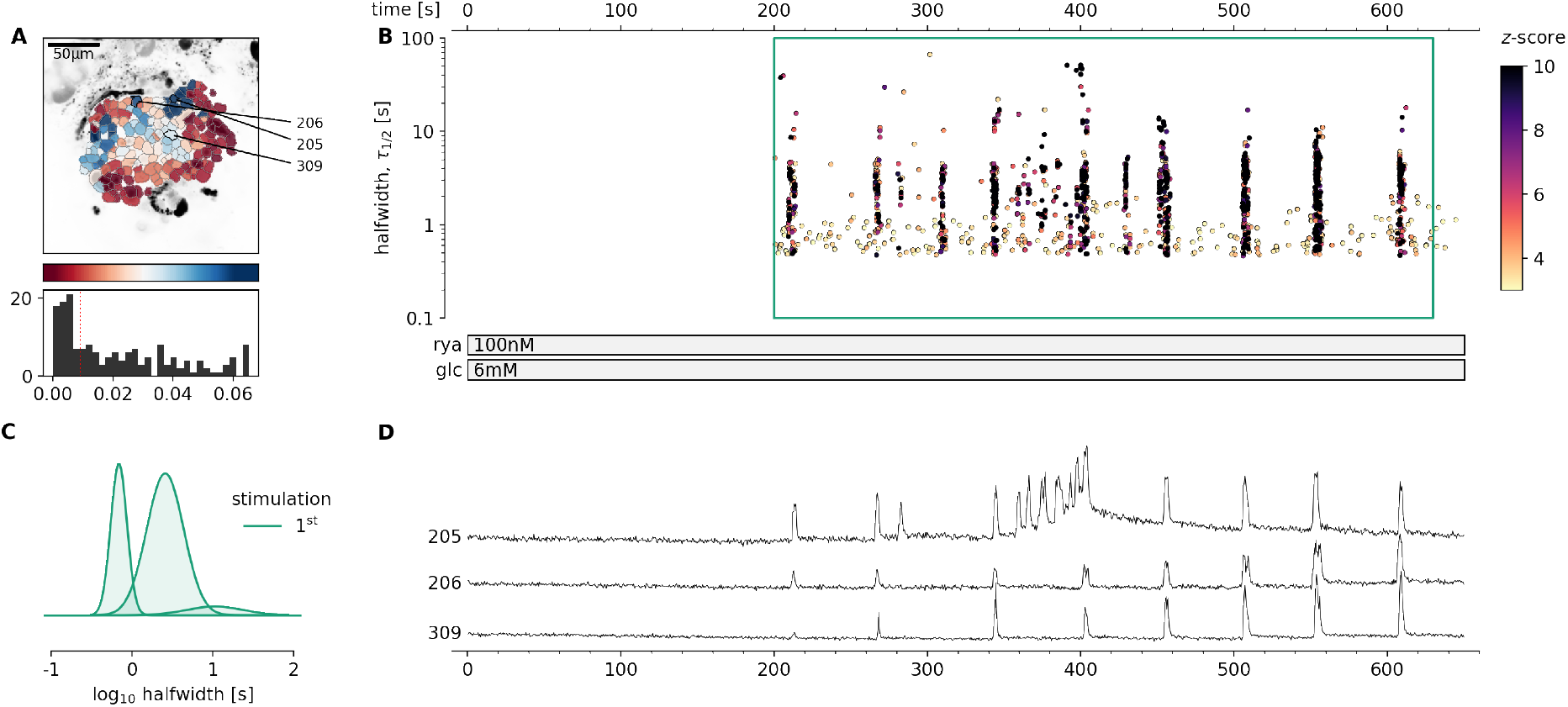
Pharmacological activation of intracellular RYR Ca^2+^ receptors in mouse beta cells at sub-threshold glucose concentration. **A**, Regions of interest (ROIs) obtained by our segmentation algorithm. The color indicates the number of events identified in the ROI trace, upon a high-pass filtering at 0.2Hz. We discarded ROIs with number of events below the threshold (red dashed line in the histogram in the lower panel). Indicated are the ROI numbers whose filtered traces correlate best with the average trace for the whole islet.**B**, Events’ halfwidth duration through time. Note the ranges of halfwidth duration occurring, the events’ synchronicity, and the phenotype’s reproducibility in ryanodine stimulation. Color indicates the statistical significance in terms of *z*-score. The stimulation protocol is indicated in the bar at the bottom of the pane. There is a prominent superposition of the short events on the plateau phases between the ROis. **C**, Normalized Gaussian fit through the logarithmic distribution of halfwidth duration during ryanodine stimulation, indicated temporal summation producing 3 discrete modes. **D**, Time courses from ROIs indicated in A, exposed to a double stimulation protocol, and rebinned to 2Hz (recorded at 20Hz). The abscissa is shared with B and is indicated there.

**Figure 3.**
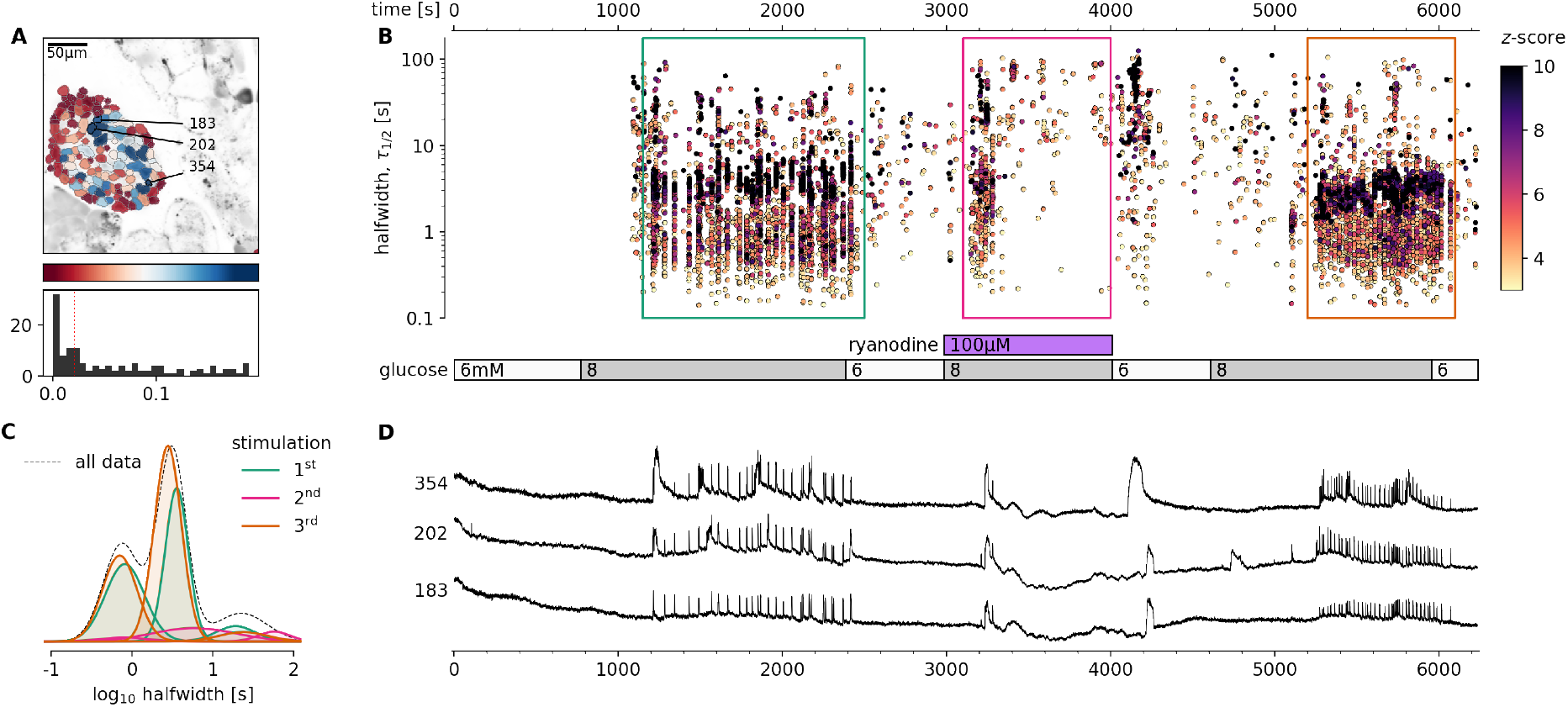
Pharmacological inhibition of intracellular RYR Ca^2+^ channels in mouse beta cells selectively inhibits the plateau [Ca^2+^]_c_ oscillations. **A**, Regions of interest (ROIs) obtained by our segmentation algorithm. The color indicates the number of events identified in the ROI trace, upon a high-pass filtering at 0.2Hz. We discarded ROIs with number of events below the threshold (red dashed line in the histogram in the lower panel). Indicated are the ROI numbers whose filtered traces correlate best with the average trace for the whole islet. **B**, Events’ halfwidth duration through time for an islet exposed to a triple 8 mM glucose stimulation protocol. Inhibitory ryanodine (100 *μ*M) was applied in the middle section of the protocol. The stimulation protocol is indicated in the bar at the bottom of the pane. There is a prominent superposition of the short events on the plateau phases between the ROis. **C**, Normalized Gaussian fit through the logarithmic distribution of halfwidth duration during control conditions and RYR inhibition, indicated temporal summation producing 3 discrete peaks. Note a complete absence of short events during the plateau phase of the second stimulation in the presence of high ryanodine. **D**, Time courses from ROIs indicated in A, exposed to a triple stimulation protocol, and rebinned to 2Hz (recorded at 20Hz). The abscissa is shared with B and is indicated there. Note the reduced [Ca^2+^]_c_ level during the exposure to high ryanodine.

The dynamic continuity of the time scales reflected in observed functional self-similarity resulted in a tri-modal distribution of halfwidth durations of the individual events that could be individually fitted with Gaussian function (Figure 1F). The reproducibility of the individual modes between different experiments was high, suggesting relatively stable lengths of unitary and compound [Ca^2+^]_c_ events (Figure 1E), and comparable degree of reproducibility and variability between the individual events on the plateau phase in the same ROI (Figure 1I), within the same ROIs during sequential stimulation with glucose (Figure 1J), within ROIs of different islet of thee same mice (Figure 1K), as well as the among islets from different mice (Figure 1L). However, the frequencies of the events of all time scales were typically not compared between different islets, since this parameter showed larger degree of variability, confirming previous reports[20]. In addition, reporting shorter modes of [Ca^2+^]_c_ dynamics in comparison to classical approaches by using low affinity indicators, we point out that these dominant short events reach at least an order of magnitude higher [Ca^2+^]_c_ and make a major contribution to Ca^2+^-dependent insulin release[26].

### Intracellular Ca^2+^ channels are sufficient to generate [Ca^2+^]_c_ oscillations at sub-stimulatory glucose

Next, we directly assessed the contribution of RYR intracellular Ca^2+^ channels, which in mouse beta cells are predominantly RYR2[16], in glucose-dependent [Ca^2+^]_c_ dynamics. We first tested the pharmacological RYR activation (Figure 2, Video 2). We showed that at sub-threshold glucose, direct stimulation with stimulatory concentrations of ryanodine (100 nM) occasionally elicited regenerative Ca^2+^ events (Figure 2). Predominantly short [Ca^2+^]_c_ events were triggered at stimulatory ryanodine concentration. The both short and long [Ca^2+^]_c_ events induced by ryanodine stimulation were a progressive temporal summation of the events in a sub-second range, and of the same duration as events observed under 8 mM glucose stimulation (Figure 1). The onset of the activity varied between individual beta cells in an islet (Figs. 2BD).

These results, together with the previously published data[27], suggest that pharmacological activation of both RYR or IP_3_ intracellular release channel could produce [Ca^2+^]_c_ events in beta cells that would be regularly observed during the 8 mM glucose stimulation. Both stimulations generated compound [Ca^2+^]_c_ events through inter-molecular CICR and presented two distinct kinetic components for intracellular release of Ca^2+^ and distinct regimes of intercellular coordination. Together these data further confirm that a selective stimulation of Ca^2+^ release from the intracellular Ca^2+^ stores can drive [Ca^2+^]_c_ oscillations in beta cells that are both coordinated (Figure 2) or non-coordinated[27].

### Intracellular Ca^2+^ channels are necessary for glucose-induced [Ca^2+^]_c_ oscillations

After demonstrating that [Ca^2+^]_c_ events can be evoked at glucose concentrations just below the glucose activation threshold with RYR activation, we used specific pharmacological tools to selectively block the activity of these intracellular channels and consequently determine the contribution of these receptors to the glucose-dependent stimulation of [Ca^2+^]_c_ events. We blocked RYR intracellular Ca^2+^ channels with an inhibitory ryanodine concentration (100 *μ*M)(Figure 3, Video 3). High ryanodine selectively and completely inhibited the dominant time scale of events and its superimposed ultra-short events during the plateau phase, leaving initial long transient events intact (Figs. 3BD). Immediately after the washout of the high ryanodine, we observed prominent long [Ca^2+^]_c_ oscillations, which is consistent with stimulatory effects of the low ryanodine concentration (Figure 3B-D). The exposure to high ryanodine concentration was fully reversible and subsequent exposure to control 8 mM glucose stimulation resulted in a response comparable to the initial control stimulation (Fig. 3B).

### [Ca^2+^]_c_ events can be initiated and maintained during L-type VACC block

Our next step towards elucidating the molecular mechanisms of the aforementioned cytosolic Ca^2+^ events was to determine to what extent these multiple time scales of [Ca^2+^]_c_ events depended on the activity of L-type VACCs. Previous experiments performed on beta cell-selective Cav1.2 Ca^2+^ channel null mice showed that, in the absence of the Ca^2+^ channel, [Ca^2+^]_c_ events were only moderately affected during the first minutes of the glucose stimulation, with a characteristic change in bursting pattern observed during acute stimulation[25]. In the present study, we used the double stimulation protocol to test the effect of a saturating concentration of isradipine (5 *μ*M), a specific L-type VACC blocker (Figure 4, Video 4). Consistent with the knockout mouse data, we observed a transient phase and initial plateau activity with a halfwidth duration with a median value of 3.1 s (Q1 1.8 s, Q3 4.6 s) during the first section and 3.4 s (Q1 2.1 s, Q3 4.3s, p=0.249) in the second section of the double protocol, where we co-applied isradipine. However, in the continuation of the plateau phase, the pattern of [Ca^2+^]_c_ events was characterised by shorter and smaller events with the median value of halfwidth duration of 2.4 s (Q1 1.9 s, Q3 3.1.s, p<10^-15^)(Figure 2C (red traces)). Instead of CICR-like compound events of the dominant time scale, a reduced number of events with a mean halfwidth duration below 2 seconds remained that could reflect progressively lower probability of CICR after blockage of VACCs and a switch towards isolated ultra-short events (Figure 4E). Within minutes after the change in the pattern of [Ca^2+^]_c_ events, the events disappeared completely.

**Figure 4.**
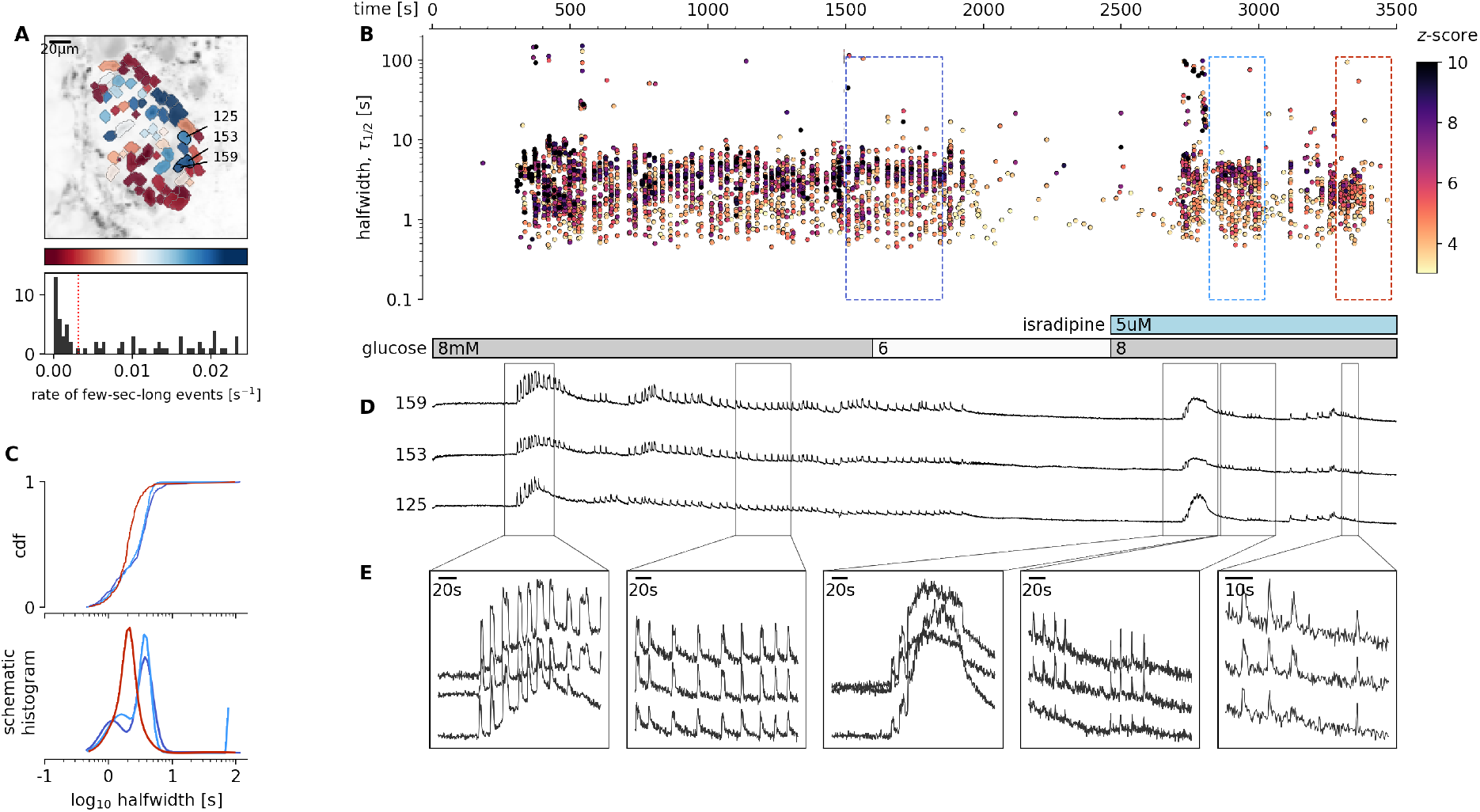
Glucose-dependent activation of beta cells in the presence of inhibitory isradipine concentration to block VACCs. **A**, Regions of interest (ROIs) obtained by our segmentation algorithm. The color indicates the number of events identified in the ROI trace, upon a high-pass filtering at 0.2Hz. We discarded ROIs with number of events below the threshold (red dashed line in the histogram in the lower panel). Indicated are the ROI numbers whose filtered traces correlate best with the average trace for the whole islet. **B**, Events’ halfwidth duration through time for an islet exposed to a double 8 mM glucose stimulation protocol. Saturating concentration of VACC blocker isradipine (5 *μ*M) was applied in the second section of the protocol. The stimulation protocol is indicated in the bar at the bottom of the pane. There is a prominent superposition of the short events on the plateau phases between the ROis. **C**, (top) Cumulative distribution frequency (cdf) of a mean halfwidth duration of events during the plateau phase of the both stimulations. (bottom) Normalized Gaussian fit through the logarithmic distribution of halfwidth duration during control conditions and inhibition of VACCs. Note a shift towards shorter events during the late plateau phase in the presence of isradipine. **D**, Time courses from ROIs indicated in A, exposed to a double stimulation protocol, and rebinned to 2Hz (original frequency is 20Hz). The abscissa is shared with B and is indicated there. **E**, Expanded time traces from a representative ROI indicating (as indicated in panel C) the a long event from the initial transient phase, followed by a plateau phase in control, long event during initiation of the second stimulation, short events from early plateau phase, and further shortened events from a late plateau phase.

Blocking L-type VACCs with isradipine during the plateau phase produced a similar change in the phenotype of [Ca^2+^]_c_ events, with progressively shorter and smaller events, before events subsided completely (Figure S4-1). A complete inhibition of the glucose-dependent [Ca^2+^]_c_ events was obtained when slices were exposed to 100 *μ*M diazoxide (Figure S4-2). At this concentration, diazoxide clamps the membrane potential close to the resting membrane potential, which is around the diffusion equilibrium potential for K^+^ (E_*K*_; −90 mV) [6, 28] and outside the range where VACCs could be activated [29]).

Our experiments confirm a role of L-type VACCs for the regenerative glucose-dependent activity of beta cells during the plateau phase[30]. However, we also confirm previous observations that these channels play only part of the role during the first minutes of beta cells activation, and that the plasma membrane depolarization plays an independent role in stimulation of intracellular Ca^2+^ release[31].

### [Ca^2+^]_c_ oscillations in low extracellular Ca^2+^ conditions

To further test the potential role of intracellular Ca^2+^ channels on ER in the spatio-temporal regulation of [Ca^2+^]_c_ events, we used a classical approach with the reduction of extracellular Ca^2+^ concentration[31]. Incubation of beta cells with 0.4 mM extracellular Ca^2+^ uncovered two major phenomena (Figure 5, Video 5). First, beta cells within the islets tended to functionally dissociate, losing coordinated and global intercellular Ca^2+^ events with a phenotype resembling the Cx36 ablated or pharmacologically blocked cell-cell electrical coupling [32, 33]. Second, low extracellular Ca^2+^ also changed the pattern of [Ca^2+^]_c_ events (Figure 5). The most prominent effect of low extracellular Ca^2+^ concentration was the complete absence of the short dominant time scale events (Figure 5), where the median halfwidth duration of 2.2 s (Q1 1.6 s, Q3 3.0 s) from the control section, decomposed to ultra-short events with a median halfwidth duration of 0.6 s (Q1 0.5 s, Q3 1.7 s, p<10^-200^) during the treatment section. In addition, the median amplitude of the events was significantly smaller in the section with low extracellular Ca^2+^ concentration. The decomposition of the dominant time scale short [Ca^2+^]_c_ events resulted in a series of smaller, but higher frequency events (Figure 5B-E), confirming previous electrophysiological measurements [6]. The coherence of responses between the individual beta cells within the islet was more localised in low extracellular Ca^2+^ concentration. Also long events were significantly longer in the low extracellular Ca^2+^ concentration with median halfwidth duration of 24 s (Q1 14 s, Q3 33 s) in comparison to 20 s (Q1 11 s, Q3 32 s, p<0.0001) during the control section. Our results suggest that in the presence of sufficient extracellular Ca^2+^ concentration, the CICR-like [Ca^2+^]_c_ oscillations are likely to occur in glucose stimulated beta cells in situ and that these events decompose at sub-physiologically low extracellular [Ca^2+^]. The spectrum of recorded patterns of beta cell activities confirms previously described role of CICR through intracellular Ca^2+^ channels in glucose-dependent activity of these cells[34, 35].

**Figure 5.**
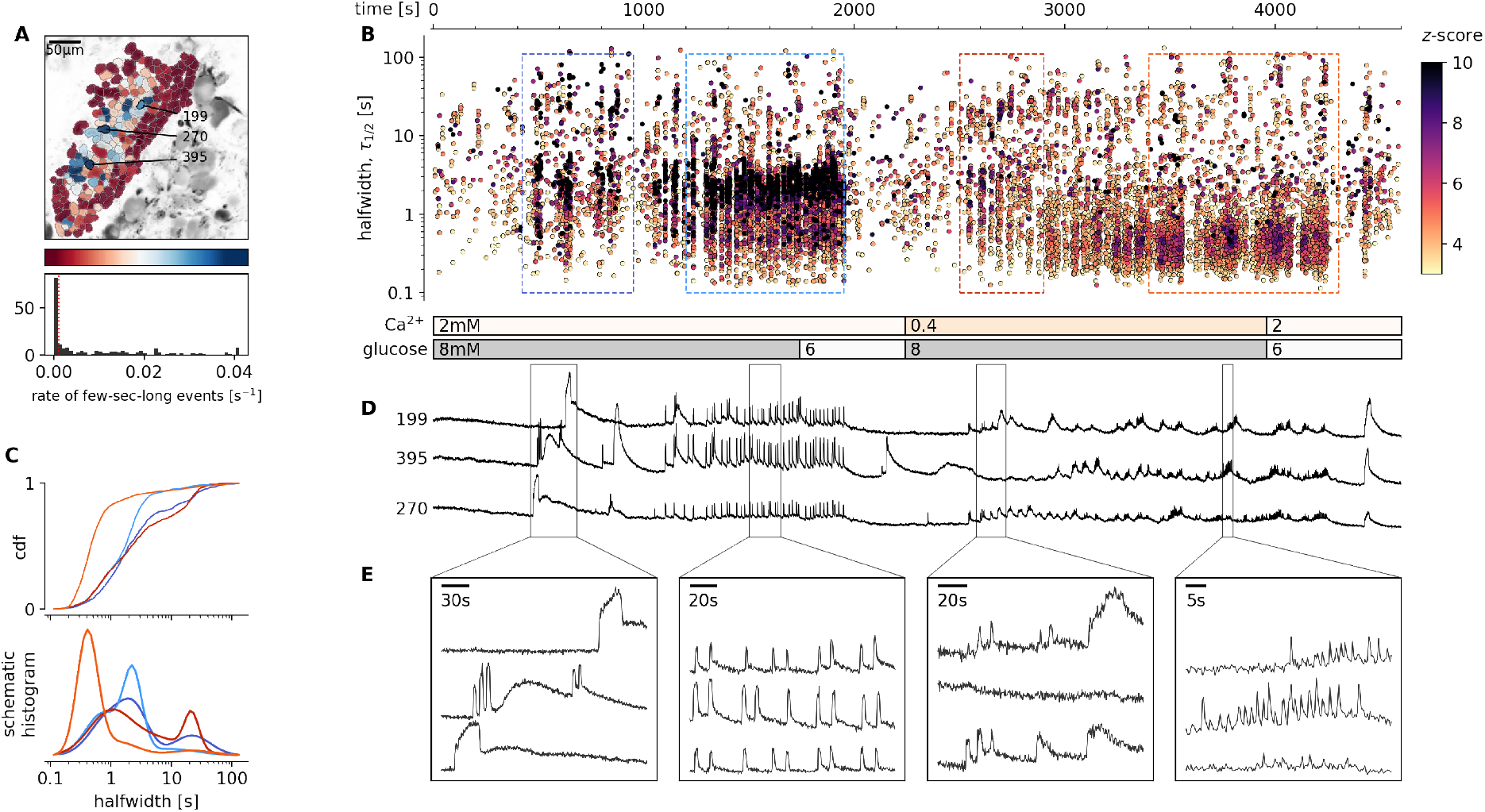
Glucose-dependent activation of beta cells at sub-physiological extracellular Ca^2+^ concentration. **A**, Regions of interest (ROIs) obtained by our segmentation algorithm. The color indicates the number of events identified in the ROI trace, upon a high-pass filtering at 0.2Hz. We discarded ROIs with number of events below the threshold (red dashed line in the histogram in the lower panel). Indicated are the ROI numbers whose filtered traces correlate best with the average trace for the whole islet. **B**, Events’ halfwidth duration through time for an islet exposed to a double 8 mM glucose stimulation protocol. Sub physiological extracellular Ca^2+^ level (400 *μ*M) was applied in the second section of the protocol. The stimulation protocol is indicated in the bar at the bottom of the pane. There is a prominent superposition of the short events on the plateau phases between the ROis. **C**, (top) Cumulative distribution frequency (cdf) of a mean halfwidth duration of events during the plateau phase of the both stimulations. Events from the initial transient phase are in dark colors, events from the plateau phase are in light color. (bottom) Normalized Gaussian fit through the logarithmic distribution of halfwidth duration during control conditions and inhibition of VACCs. Note a shift towards shorter events during the plateau phase in the conditions of sub physiological extracellular Ca^2+^ concentration.**D**, Time courses from ROIs indicated in A, exposed to a double stimulation protocol, and rebinned to 2Hz (recorded at 20Hz). The abscissa is shared with B and is indicated there. **E**, Expanded time traces from a representative ROI indicating (as indicated in panel C) the a long event from the initial transient phase, followed by a plateau phase in control, long event during initiation of the second stimulation, and short events from early plateau phase in low extracellular Ca^2+^ concentration.

The low extracellular Ca^2+^ concentration recordings provide further evidence regarding the existence of two distinct phases for the activation and activity of beta cells. The initial phase during the activation of beta cells in the fresh pancreas slice consisted of one or few long IP_3_-dependent [Ca^2+^]_c_ transients (some tens of seconds long) as reported before[27], followed by a plateau phase comprising of a series of short RYR-mediated events.

## DISCUSSION

Decades of electrophysiological experiments on pancreatic beta cells have established a critical role for plasma membrane ion channels in controlling excitability, dynamics of cytosolic Ca^2+^ events and insulin exocytosis [3, 23]. In standard electrophysiology experiments, beta cells are kept at 3 mM glucose where they are electrically silent, after which they are activated by an instantaneous increase to glucose levels well in excess of 10 mM [20, 33].

In the current study, we combined fast confocal microscopy, a photostable and bright low affinity Ca^2+^ sensor, and tools of data sciences to obtain new insights into physiological activation of beta cells within intact islets in fresh pancreas tissue slices. Specific pharmacological modulation of RYR in intact beta cell collectives stimulated with 8 mM glucose and recorded with a unique spatio-temporal resolution demands an updated model of beta cell activation and bursting activity. According to our model, multiple time scales of events are generated with a progressive temporal superposition of ultra-short events, which represent CICR of either IP_3_[27] or RYR Ca^2+^ release channels. Both intracellular channels can be directly stimulated by glucose. The ultra-short events represent unitary activity of intracellular Ca^2+^ release channels, which can engage in intra-molecular CIC, and as described for other cell types, depend on sufficient Ca^2+^ ER load[36, 37].

Evidence for CICR in beta cells involving both IP_3_ and RYR channels has been previously reported [38–40]. When, however, the ER Ca^2+^ load decreases, CICR bursts switch to sub-second events produced by individual or more localized clusters of intracellular Ca^2+^ channels before events eventually stop. Decomposed CICR bursts therefore appear to oscillate at higher frequency, a phenomenon reported after ER emptying using thapsigargin [41], genetic ablation of L-type VACCs [25], or lowered extracellular Ca^2+^[6, 41]. Our experiments show that physiological glucose concentrations (e.g. 8 mM) support sufficient ER Ca^2+^ load [42] to enable CICR bursts triggered by direct pharmacological stimulation of RYR and IP_3_ Ca^2+^ channels. Addition of supra-threshold glucose concentration first activates a prominent IP_3_-dependent transient Ca^2+^ release lasting for several tens of seconds[27]. IP_3_ activation is localized to segregated beta cell clusters and not coordinated within the whole islet[43]. Next, Ca^2+^-dependent mitochondrial metabolism of glucose boosts ATP production[15], that directly decreases the opening probability of K_*ATP*_ channels[44], and directly activates RYR Ca^2+^ channels[45, 46]. High input resistance due to K_*ATP*_ channel closure enhances the coordination between beta cells during the regenerative RYR-CICR activity. We also observed coherent intercellular activity after ryanodine stimulation at sub-stimulatory glucose and this activity was specifically blocked by inhibitory concentrations of ryanodine, suggesting that RYR independently promotes intercellular coordination due to strategic cellular localisation of these receptors. The long-term activity of intracellular Ca^2+^ receptors during the plateau phase of beta cells must be supported with mobilisation of Ca^2+^ from the extracellular space, where VACCs could activate through a large ER Ca^2+^ depletion as has been suggested earlier[41]. Our results do not discount a role of plasma membrane K_*ATP*_ channels and voltage-gated Ca^2+^ entry in beta cell function. We speculate that K_*ATP*_ channels play a critical depolarizing role to support activation of L-type VACCs, reloading the ER Ca^2+^ stores, promoting cell-cell coordination and the long-term regenerative activity, consistent with the human genetics of beta cell responsiveness and their clinical utility as targets of sulphonylurea therapy[47]. K
_*ATP*_ channels may also play an outsized role in the response to an instantaneous glucose increase from hypoglycemic levels (2-3 mM) to supra-physiological levels (15-25 mM). This is consistent with previous observations suggesting closure of K_*ATP*_ channels is not the sole mechanism to depolarise beta cells[48]. Indeed, other plasma membrane ionic currents have been invoked to explain the complex oscillatory behaviour of beta cells [3, 23]. As an example, multiple Ca^2+^-dependent K^+^ and other K^+^ channel activities have been bundled to explain the elusive, so called K_*slow*_ conductance, that could critically shape the typical electrical bursting pattern of beta cells after initial beta cell activation [49–52]. The details about the physiological glucose responses are even less clear in human beta cells and still need to be performed[53].

Glucose plays an essential role in filling, and therefore also emptying, intracellular Ca^2+^ stores[42]. Forced depletion of intracellular Ca^2+^ stores using thapsigargin or extracellular EGTA, increases cytosolic Ca^2+^ concentration, and produces a sustained depolarisation, with increased frequency of membrane potential oscillations[6, 41]. Similarly, reduced activity of the SERCA2 pump to maintain ER Ca^2+^ loading has been shown to disrupt glucose-stimulated calcium signalling[54]. Increased frequency of [Ca^2+^]_c_ events was also recorded in our experiment using low extracellular Ca^2+^ concentration. We demonstrate that glucose around the threshold (6-7 mM glucose) supports ER Ca^2+^ release through IP_3_[27] and RYR release channels. In addition to support sufficient Ca^2+^ load in the ER, glucose-dependent effects in beta cells provide all key substrates, like ATP and cAMP, to directly trigger and modulate the activation of intracellular Ca^2+^ release channels, even in the absence of plasma membrane depolarization that would increase the opening probability of VACCs. Included in these stimuli is the parasympathetic release of ACh binding to muscarinic ACh receptors (mAChRs) and induce insulin release via the production of IP_3_ and Ca^2+^ release from the intracellular stores[27, 55–57]. Previous studies, including our own, systematically underestimated the role of RYR and IP_3_ receptors due to pre-emptying of intracellular Ca^2+^ stores in too low (0-3 mM) extracellular glucose.

The prominent role for intracellular Ca^2+^ release had strong early support from ^45^Ca^2+^ flux studies[30], but subsequent electrophysiological work challenged this paradigm[41, 45] and evidence accumulated in favor of the dominance of plasma membrane K^+^ channels and VACCs in patterning the dynamics of [Ca^2+^]_c_ events. Even the presence of RYR Ca^2+^ channels in beta cells was debated[32, 58]. There has been less controversy regarding the expression and roles of IP_3_Rs [59, 60]. However, we and others have documented RYR activity and confirmed RYR2 as the most abundant isoform expressed in rat, mouse, and human beta cells[13, 14, 16, 61, 62]. Indeed, the unique localization and Ca^2+^-release kinetics of each intracellular Ca^2+^ release channel enables the coding of Ca^2+^ signals to control specific cellular[62–65] or intercelllar functions. The duality of contributions of both IP_3_ and ryanodine receptors has been previously described to underly complex patterns of intracellular Ca^2+^ regulation of neuronal activity[66]. As proposed in our recent theoretical paper, beta cell and islet collectives could utilize dual Ca^2+^ sources to adequately respond to a variety of metabolic challenges from those requiring transient hormonal adjustments to those requiring major release of insulin[67]. Such a functional arrangement would support both a form of a circuit memory, and risk of a stochastic editing contributing to the pathogenesis of diabetes mellitus[68].

It was proposed that a beta cell’s ER is extraordinarily leaky to Ca^2+^ [69]. ER Ca^2+^ leak is accelerated by ER stress, leading to beta cell apoptosis[14, 16]. In addition, excitotoxicity and ER Ca^2+^ overload have been implicated in beta cell apoptosis [16] and may also contribute to diabetes pathogenesis. There is strong evidence that ER dysfunction is involved in the pathogenesis of both type 1 and type 2 diabetes [70]. In type 1 diabetes, ER dysfunction is a prominent and early feature [71]. In type 2 diabetes, Genome-wide association studies have identified multiple RYR2 SNPs with suggestive evidence in dozens of glycemic traits (www.type2diabetesgenetics.org, accessed Feb 2021)[72].

Both ER Ca^2+^ load as well as intracellular Ca^2+^ channels can serve as targets for therapy of diabetes mellitus[73]. Pharmacological inhibition of VACCs activity with verapamil was recently reported to have a positive effect on beta cell function and survival in adults with recently diagnosed T1D [74], consistent with previous pre-clinical studies [75]. Pre-clinical studies also demonstrate that inhibition of voltage-gated Na^+^ channels can protect beta cells from cytokine-induced death, and reduces diabetes incidence in NOD mice[76, 77]. Also, intracellular Ca^2+^ release channels are druggable and were implicated in successful diabetes therapies. For example, RYR2 receptors can be activated by both ATP and PKA [46] and, can therefore be directly stimulated by glucose and modulated by incretins, catecholamines, and peptide hormones. RYRs are a proposed therapeutic target in Wolfram syndrome[78]. Abundant physiological data also indicate that CICR from intracellular stores can serve as an “amplifier” of glucose-induced insulin granule exocytosis and plays a central role in incretin-induced insulin secretion[34, 79].

The major strength and limitations of our approach are essentially the same as they have been described for a fresh pancreatic slice preparation, which we introduced in 2001[18]. This is one of the first studies where we could use high spatial and temporal resolution imaging to specifically address intracellular Ca^2+^ receptors. Some of the observed long events may result from phenomena not necessarily connected to insulin secretion, such as movement, cilia protrusion, metabolism, drift of the objective of the microscope. At the moment we do not have a full understanding of the direction and the magnitude of all potential biases beyond the presented experimental evidence. In summary, using powerful new rapid imaging and data analysis methods, we show here that RYR play essential roles in glucose stimulated Ca^2+^ signalling in beta cells in situ.

## MATERIALS AND METHODS

### Ethics statement

We conducted the study in strict accordance with all national and European recommendations on care and handling experimental animals, and all efforts were made to minimise the suffering of animals. The Ministry of Education, Science and Research, Republic of Austria (No: 2020-0.488.800) and administration of the Republic of Slovenia for Food Safety, Veterinary and Plant Protection (No: U34401-12/2015/3) approved the experimental protocol (No: U34401-12/2015/3).

### Tissue slice preparation and dye loading

C57BL/6J mice, 8–20 weeks of age, and of either sex (Jackson Laboratories), were kept on a 12:12 hours light: dark schedule in individually ventilated cages (Allentown LLC, USA) and used to prepare pancreatic tissue slices, as described previously [8, 80]. In brief, after sacrificing the mice, we accessed the abdominal cavity via laparotomy and distally clamped the common bile duct at the major duodenal papilla. Proximally, we injected the low-melting-point 1.9 % agarose (Lonza, USA) dissolved in extracellular solution (ECS, consisting of (in mM) 125 NaCl, 26 NaHCO_3_, 6 glucose, 6 lactic acid, 3 myo-inositol, 2.5 KCl, 2 Na-pyruvate, 2 CaCl_2_, 1.25 NaH_2_PO_4_, 1 MgCl_2_, 0.25 ascorbic acid) at 40 °C into the common bile duct. Immediately after injection, we cooled the agarose infused pancreas with ice-cold ECS and extracted it. We prepared tissue slices with a thickness of 140 *μ*m with a vibratome (VT 1000 S, Leica) and collected them in HEPES-buffered ECS at RT (HEPES-buffered ECS, consisting of (in mM) 125 NaCl, 10 HEPES, 10 NaHCO_3_, 6 glucose, 6 lactic acid, 3 myo-inositol, 2.5 KCl, 2 Na-pyruvate, 2 CaCl_2_, 1.25 NaH_2_PO_4_, 1 MgCl_2_, 0.25 ascorbic acid; titrated to pH=7.4 using 1 M NaOH). For staining, we incubated the slices for 50 minutes at RT in the dye-loading solution (6 *μ*M Calbryte 520, AAT Bioquest), 0.03 % Pluronic F-127 (w/v), and 0.12 % dimethylsulphoxide (v/v) dissolved in HEPES-buffered ECS). The Ca^2+^ fluorescent dyes from a Calbryte series are highly fluorescent and photostable, allowing long-term recording even in deeper layers of the tissue slice. The linear part of the Ca^2+^-binding curve for Calbryte 520 (KD of 1.2 *μ*M) captures the [Ca^2+^]_c_ changes in beta cells better than the high affinity sensors we routinely used in our imaging experiments before. All chemicals were obtained from Sigma-Aldrich (St. Louis, Missouri, USA) unless otherwise specified.

### Stimulation protocol and cytosolic calcium imaging

We transferred individual pancreatic slices to a perifusion system containing 6 mM glucose in HEPES-buffered ECS at 34 °C. We show representative traces for each experimental condition. Each of these conditions has been tested on at least 3 separate experimental days. Slices, with the exception of experiments presented in Figs. 4 and 5, were exposed to a series of three square-pulse-like stimulations (triple protocol) characterised by exposure to 8 mM glucose for 25 minutes, followed by a washout with sub-stimulatory 6 mM glucose concentration until all the activity switched off. We typically applied the pharmacological treatment during the second section of the triple protocol (section #2). Imaging was performed on standard confocal microscopes equipped with resonant scanners (Leica Microsystems TCS SP5 and SP8 or Nikon A1R) both upright and inverted with their respective sets of 20x high NA objective lenses. Acquisition frequency was set to at least 20 Hz at 256 x 256 pixels, with pixels size close to 1 *μ*m^2^ to allow for a precise quantification of [Ca^2+^]_c_ oscillations. Calbryte 520 was excited by a 488 nm argon laser (Leica SP5 and Nikon A1R) or 490 nm line of a white laser (Leica SP8). The emitted fluorescence was detected by HyD hybrid detector in the range of 500-700 nm using a photon counting mode (Leica) or GaAsP PMT detectors (Nikon).

### Analysis and processing of data

The general analysis pipeline was as follows. Experiments involving imaging of pancreatic slices typically focused on a single field of view showing up to hundreds of cells, in a recording of at least several, often dozens, gigabytes. Current tools that are widely used (e.g., ImageJ) rely on loading the recording, or its part, into memory, for viewing, analysis, and processing. It also requires laborious and long human engagement. We have developed a set of interdependent tools to automatise as much as possible the analysis pipeline (Figure 6, Supporting information).

**Figure 6.**
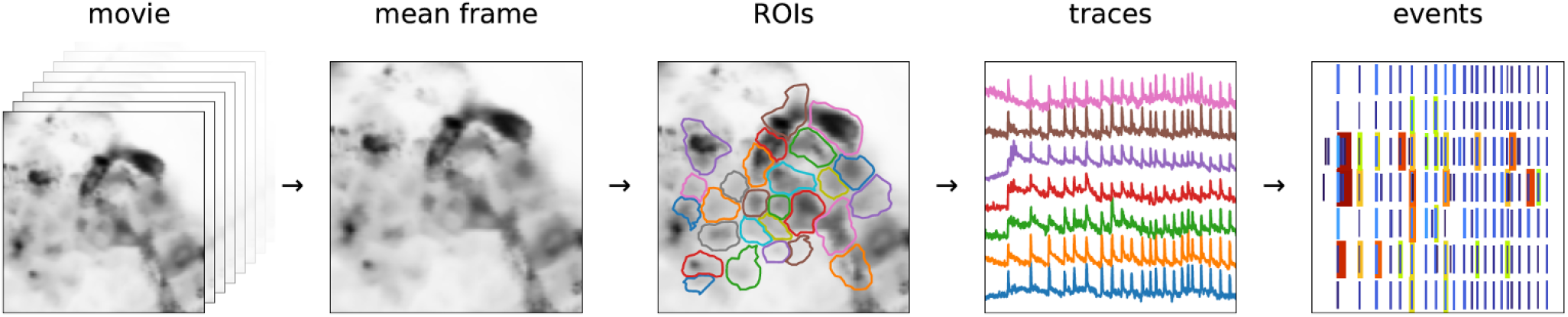
Processing pipeline to automatically detect ROIs and [Ca^2+^]_c_ events at all time scales within an experiment. From a full movie, we calculated the mean (or other statistic) across all frames. We passed the mean image through a band-pass filter and define ROIs by detecting local peaks of light intensity. We then saved ROIs with all the important information (time traces, ROI coordinates, movie statistics, recording frequency, pixel size, etc.). Traces contained features at very different timescales - with different timescales presumably being important for different cell types. We collected them into separable events for analysis.

Regions of interest (ROIs) were identified using a semi-automatic detection as follows. Recordings were stored as a 3-dimensional (*T*×*X*×*Y*) numpy array [81]. When the recording was stable, obtaining a mean image, or any other statistic over frame, was rather trivial. In case there was horizontal movement, it could be corrected for by aligning the frames to a template. For this we used the functionality present in CaImAn [82], except that high frequency recordings needed to be rebinned to some moderate frequency (a few Hz), before correcting, in order to reduce the noise level. Once the translation offsets were obtained, we used them to correct the recording in original frequency. To define regions of interest, we blurred the representative image by a kernel of the size we expect cells to be, and at the scale double of that. The difference between these two images represents a band-pass filter of the original, where the local intensity variations were emphasised (Figure S6-1). We then passed through all pixels where the value of the filtered image was positive (or larger than a small positive threshold), and for each pixel we searched for a local peak in its vicinity. All the pixels that lead to the same local peak were then grouped into a single ROI. As we were mainly interested in islet cells, we chose the kernel size to approximately correspond to 10 *μ*m, which is the characteristic length scale of the islet cells. If the pixel size is unknown, the choice of kernel is up to the person running the scripts.

Representative image can be a mean over all frames or any other statistic. In addition, our code supports standard deviation, mean and standard deviation of the first derivative of the movie, and a “robust maximum” of the movie. As “robust maximum”, we defined a very high percentile of the set absolute values of a set, essentially a value close to its maximum, by default it is 10th largest. We avoid the maximum as a means to make analysis robust to outliers. This statistic is sensitive to cells which fire extremely rarely during a recording, so that the mean of those pixels is negligible. By default, we choose an average of the mean and high percentile as a representative image for band-pass filtering and ROI extraction.

Trace processing was conducted as follows. In an ideal detector, recording a time trace of a single pixel in absence of any signal would consist of independent values of the number of photons detected during the dwell time. The values (*x*) are then distributed according to the Poisson distribution, with standard deviation (*σ*_1_) being equal to the square root of the mean 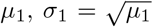. Transformation to standard score or *z*-score (Figure S6-2) with 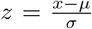 is then performed that recasts the initial quantity x in the units of standard deviation from the expected mean.

A noisy trace in *z* spends 95% of the time between −2 and 2, and 99.7% between −3 and 3. Probability of *z* > 3 is very small *p* < 0.0013, which is why it is often considered a threshold value for pointing out the outliers. In general, the mean slow component needed to be inferred, typically by low-pass filtering. Throughout this project, we used cascaded second-order section (sos) filtering, implemented in scipy.signal module [82]. The cut-off frequency *f_cut_* of a filter determines the timescale of the events that can be detected in *z*-score.

Fast Fourier transform naturally distorts signals, but the inferred *z*-score can be used to correct for it. We constructed an iterative filtering algorithm, where at each iteration, we neglect the outlier values of the original trace, substitute them with the values of the slow component, and reapply the filter. At each iteration, the distortion is less prominent, increasing the *z*-score. In Fig. S6-2, we show the result after 10 iterations, though we found three iterations as a more conservative choice, and a reasonable compromise between results and computing time.

All above refers also to a sum of pixel traces, but, crucially, not to their mean. A sum of two pixels *a* and *b* (*x* = *x_a_* + *x_b_*), with means *μ_a_* and *μ_b_*, would have a standard deviation as expected 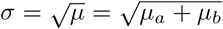. But, if we were to consider the average 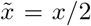, standard deviation would be 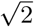 times smaller 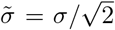. Therefore, when calculating *z*-score for a ROI trace, we always considered the sum, rather than the average trace of the underlying pixels. When we visualised traces, we show them averaged, only to keep the scales comparable. The same reasoning holds for rebinning a single trace, where the resulting trace, rebinned by a factor of *n*, had a standard deviation 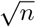 times smaller than the original.

Experiments discussed in this manuscript were recorded on standard Leica SP5 or SP8, as well as NIKON A1R confocal microscopes. In cases where we used a Hybrid detector in the photon counting mode (Leica) we saw no significant departure from our assumption of Poisson distribution. Even with non-unit gain, the linear dependence between variance and mean remains, though the slope was different from 1 (Figure S6-3). Other types of detectors introduce additional sources of noise other than Poisson (eg. thermal), but mostly they were still dominated by Poisson in our experience, at least as long as the values between neighbouring pixels and frames were independent.

Traces contain features spanning orders of magnitude in time: from tens of milliseconds, to tens of minutes. We aimed to investigate how these events at different timescales and a connection between them and the islets’ environment interact. For this, we devised a two-step algorithm to identify intrinsic timescale of events and to minimise false positives (Figure S6-4). In the first step, we performed a sequential filtering of the traces at timescales starting from 0.5s, and increasing by a factor of 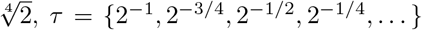, until the timescale of the longest event of interest was achieved. At each timescale, we transformed the trace to *z*-score, and identify regions where *z* > 4 as *candidate events*.

Events were characterised by the start time ((*t*_0_), its maximal height, and the width at the half of the height (*halfwidth, δt*), which is our measurement of its duration. For simplicity, we defined end time as *t*_end_ = *t*_0_ + *δt*, although events arguably last much longer after the intensity drops at half of the peak.

For an event to be considered real, it needed to be detected at multiple timescales, and will have started around the same time and will have approximately the same halfwidth. We specify a tolerance of 20 % of the halfwidth as to whether two candidate events should be considered equal; if their start and end times were within 20 % of the halfwidth, they were considered cognates, having arisen due to the same real event. For a set of cognates, we estimated the start and end time of the real underlying event as a median over the set. If the resulting estimate for the halfwidth is larger than 2 s, we required that a set consists of at least 4 candidate events (corresponding to event being detectable when filtered at timescales that differ at least two-fold). For shorter events, we required only that an event is not unique to a single timescale.

We also neglected events that last less than 3 time-frames, as well as those too close to the beginning or end of the recording (within *δt*/2), which we ascribed to artefacts from zero-padding for the filtering. We termed this procedure event *distilling* (Figs. 6, S6-4).

## ACKNOWLEDGEMENTS

MSR received grants by the Austrian Science Fund/Fonds zur Förderung der Wissenschaftlichen Forschung (bilateral grants I3562-B27 and I4319-B30). MSR, CE-M and JDJ received financial support from NIH (R01DK127236). MSR, AS and DK further received financial support from the Slovenian Research Agency (research core funding program P3-0396 and projects N3-0048, N3-0133 and J3-9289). JDJ receives grants from CIHR and Diabetes Canada.

## SUPPORTING INFORMATION

### SUPPORTING VIDEOS

Link to videos: https://figshare.com/s/e6d3945b658ec4ec3989

**Video** Figure 1.

**Video** Figure 2.

**Video** Figure 3.

**Video** Figure 4.

**Video** Figure 5.

**Figure S4-1.**
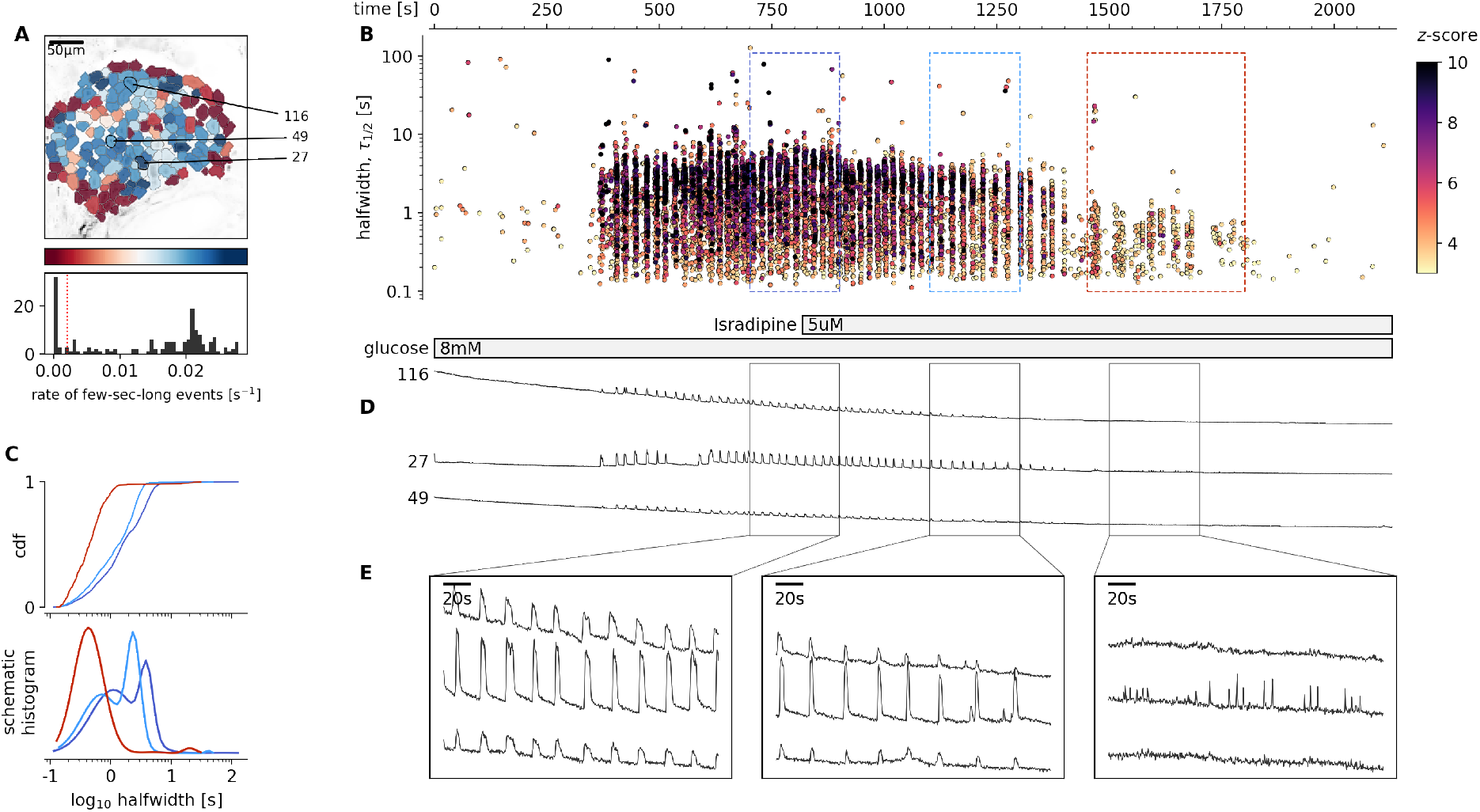
Glucose-dependent plateau activity of beta cells after blockage of VACCs. **A**, Regions of interest (ROIs) obtained by our segmentation algorithm. The color indicates the number of events identified in the ROI trace, upon a high-pass filtering at 0.2Hz. We discarded ROIs with number of events below the threshold (red dashed line in the histogram in the lower panel). Indicated are the ROI numbers whose filtered traces correlate best with the average trace for the whole islet. **B**, Events’ halfwidth duration through time for an islet exposed to a single 8 mM glucose stimulation protocol. Saturating concentration of isradipine (5 *μ*M) was applied during the plateau phase of the activity. The stimulation protocol is indicated in the bar at the bottom of the pane. There is a prominent superposition of the short events on the plateau phases between the ROis. **C**, (top) Cumulative distribution frequency (cdf) of a mean halfwidth duration of events during the plateau phase of the both stimulations. Events from the initial transient phase are in colored as indicated in panel B. (bottom) Normalized Gaussian fit through the logarithmic distribution of halfwidth duration during control conditions and inhibition of VACCs. Note a shift towards shorter events during the plateau phase in the presence of isradipine. (**D**) Time courses from ROIs indicated in A, exposed to a single stimulation protocol, and rebinned to 2Hz (recorded at 20Hz). The abscissa is shared with B and is indicated there. **E**, Expanded time traces from a representative ROI indicating (as in panel C) events from the different sections of the plateau phase.

**Figure S4-2.**
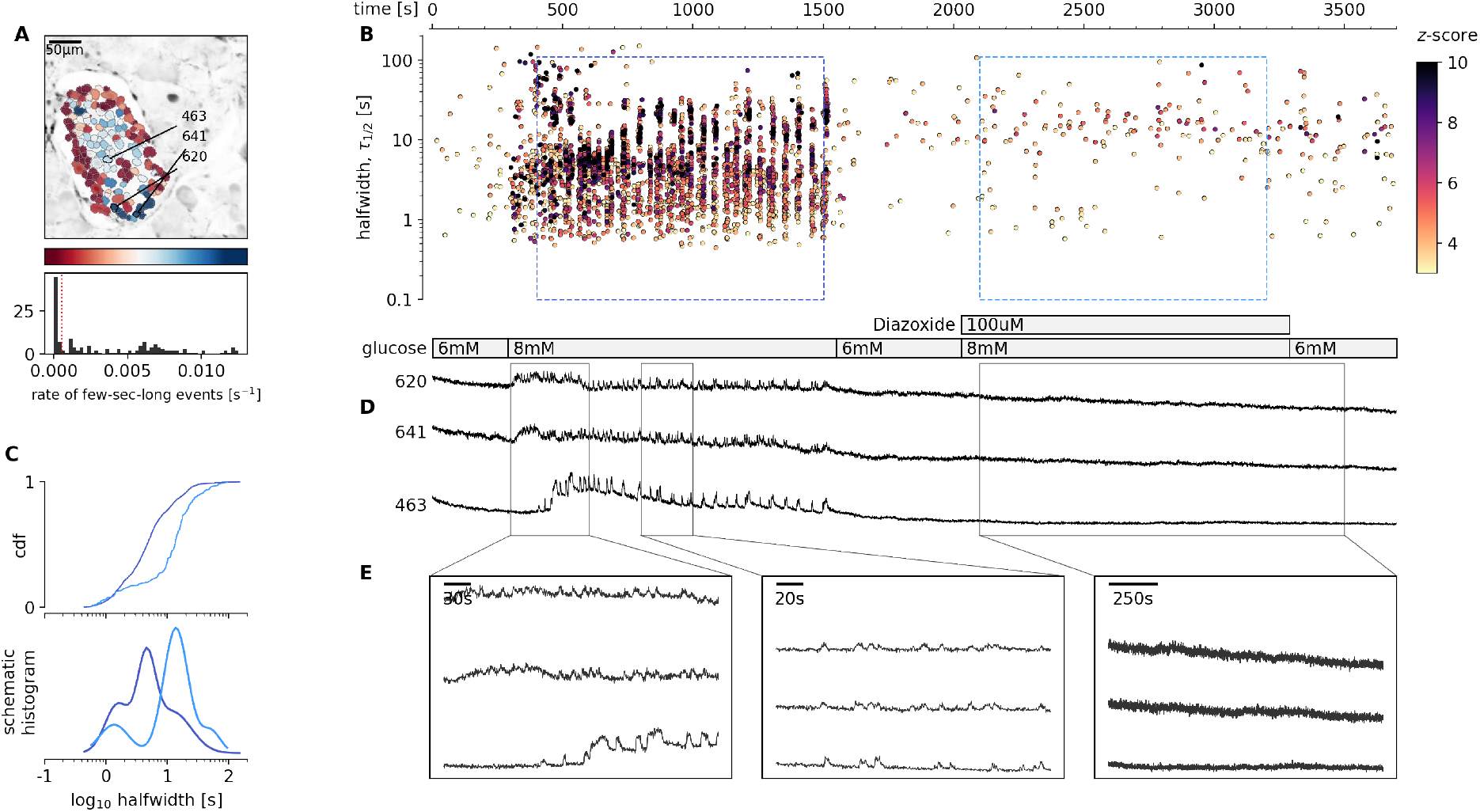
Glucose-dependent activation of beta cells in the absence of the plasma membrane depolarisation. **A**, Regions of interest (ROIs) obtained by our segmentation algorithm. The color indicates the number of events identified in the ROI trace, upon a high-pass filtering at 0.2Hz. We discarded ROIs with number of events below the threshold (red dashed line in the histogram in the lower panel). Indicated are the ROI numbers whose filtered traces correlate best with the average trace for the whole islet. **B**, Events’ halfwidth duration through time for an islet exposed to a single 8 mM glucose stimulation protocol. Saturating concentration of isradipine (5 *μ*M) was applied during the plateau phase of the activity. The stimulation protocol is indicated in the bar at the bottom of the pane. There is a prominent superposition of the short events on the plateau phases between the ROis, exposed to a double 8 mM glucose stimulation protocol. 100 *μ*M diazoxide was applied in the second section of the protocol. **C**, (top) Cumulative distribution frequency (cdf) of a mean halfwidth duration of events during the plateau phase of the both stimulations. Events from the first phase are in dark color, events from the second plateau phase are in light color. (bottom) Normalized Gaussian fit through the logarithmic distribution of halfwidth duration during control conditions and in the absence of plasma membrane depolarization. Note a shift towards longer events during the plateau phase in the conditions of high concentration of diazoxide. (**D**) Time courses from ROIs indicated in A, exposed to a double stimulation protocol, and rebinned to 2Hz (recorded at 20Hz). The abscissa is shared with B and is indicated there. **E**, Expanded time traces from a representative ROI indicating (as in panel C) events from the different sections of the plateau phase.

**Figure S6-1.**
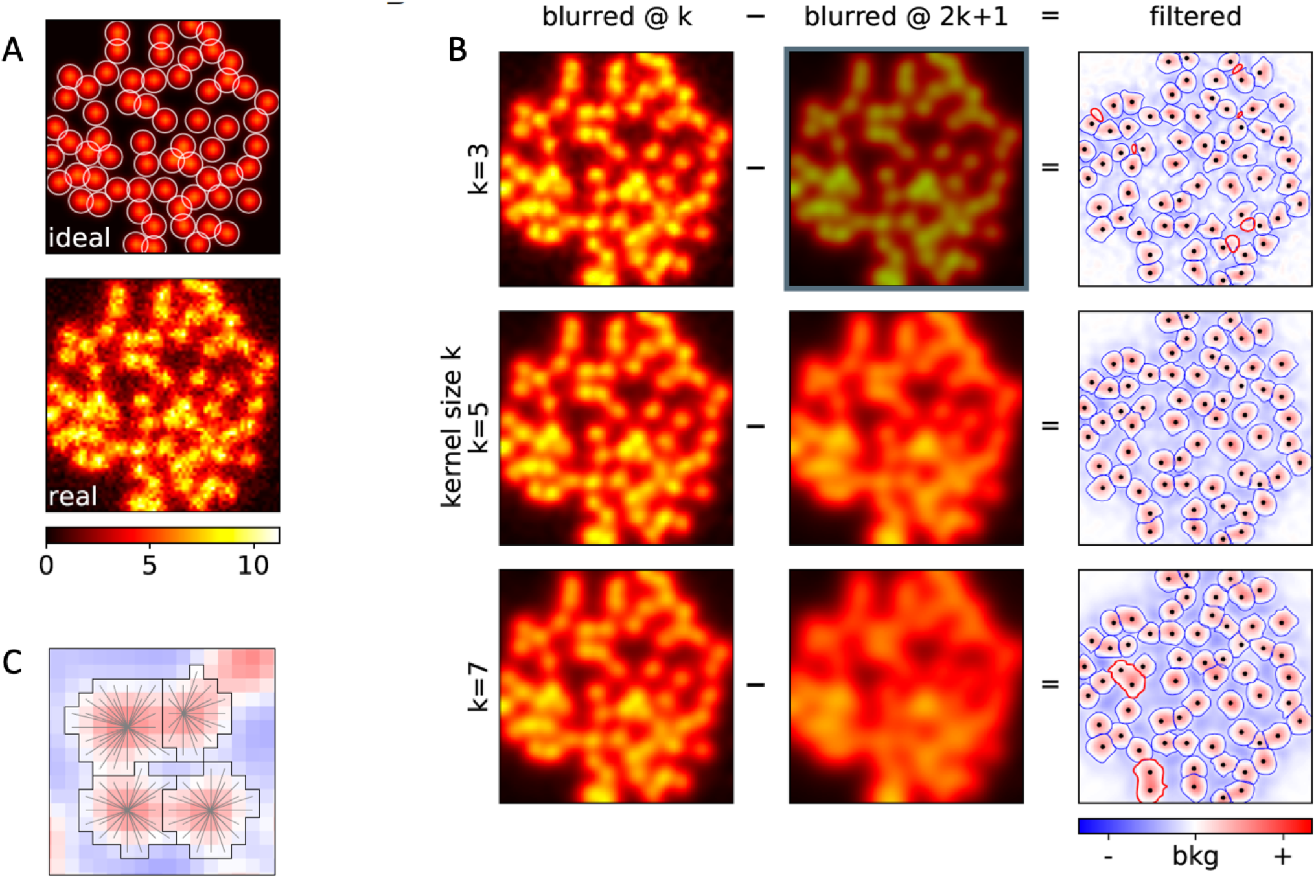
**A**, Computationally created template 64×64 image with uniform spherical cells, and with Poisson noise added. **B**, Band-pass filtering of the “realistic” image from A, with different kernel sizes, for denoising (*left*), and for approximating background (*middle*). The difference is the filtered image, used to construct ROIs (*right*). The size of the kernel determines the approximate size of the ROIs obtained. In thick red contours we emphasise misidentified ROIs; the dots indicate the real locations of the cells’ centers. **C**, Each ROI is constructed by explicitly searching for a closest peak in intensity. A pixel can only be part of a single ROI.

**Figure S6-2.**
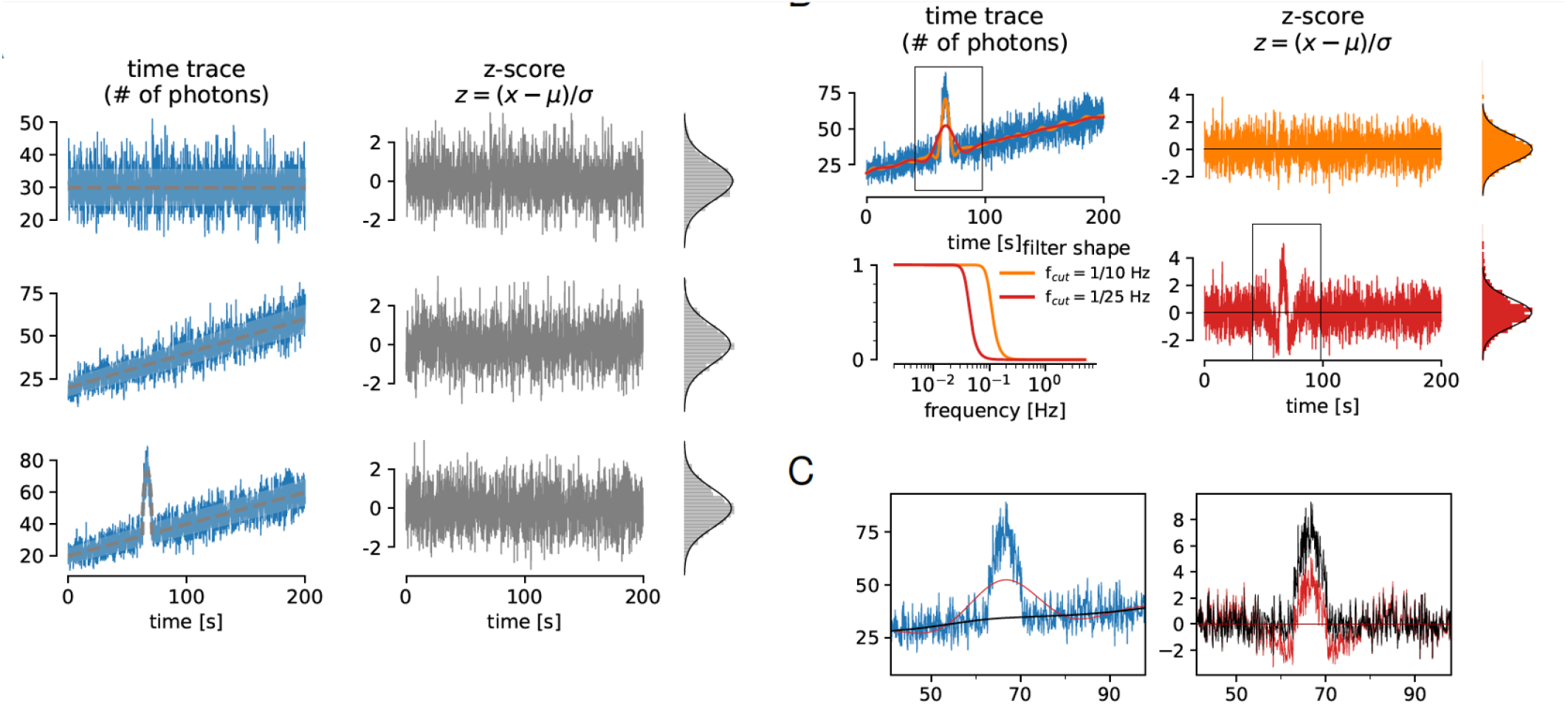
**A**, *Left*: Simulated traces with Poisson noise (blue) around different means (grey) with different temporal features. The shaded area represents one standard deviation interval around mean. *Right*: Irrespective of the shape of the mean, *z*-score transformation behaves exactly as a Gaussian variable with zero mean and unit standard deviation. **B**, The values of *z*-scores depend on the reference. In subfigure (A), the reference curves are known, but in general they need to be inferred, typically by low-pass filtering. The cut-off frequency *f_cut_* of a filter determines the timescale of the events that can be detected in *z*-score. If filtered with too high cut-off (orange), the resulting smooth curve follows too closely the trace and the event is not visible in *z*. With *f_cut_* = 1/50 Hz, the resulting *z*-score has obvious outliers (*z* ≥ 3), visible both in trace and in the histogram. **C**, Zoom to the indicated regions form B. The original (blue) and the filtered trace and *z*-score (red), and the filtered trace and *z*-score corrected for distortion (black).

**Figure S6-3.**
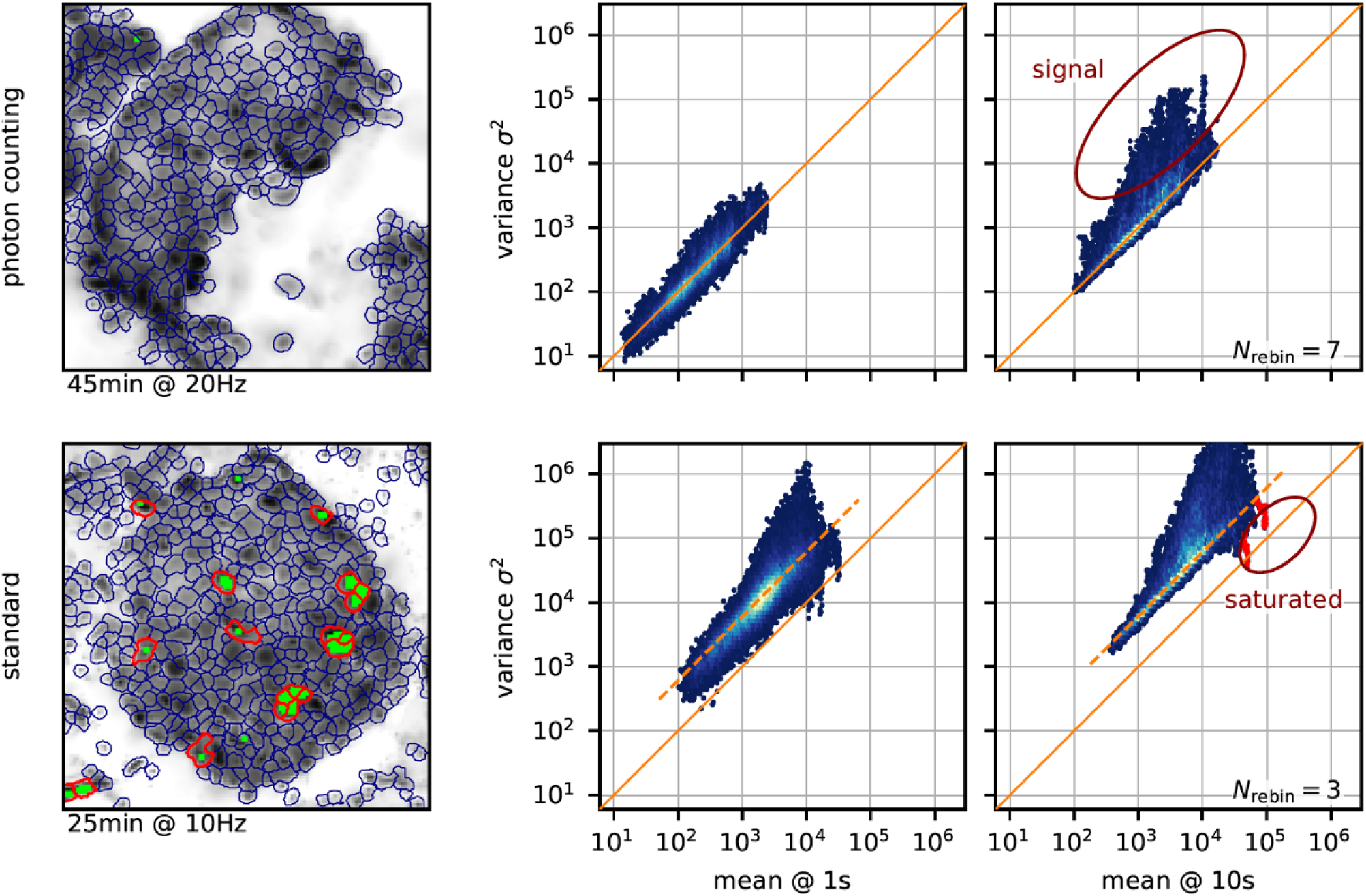
*Left*: Regions of interest for the two experiments recorded using the raw photon-counting mode (*top* and the standard mode with large gain (*bottom*). Pixels which had at least 10 saturated values are shown in green. In thick red contours we emphasise ROIs with at least 30 green pixels used to emphasise the influence of saturation (see below). For each recording, we filter the ROI traces at 1s (*middle*) and 10s (*right*) to separate the trace into slow and fast component. For each ROI we then randomly choose 100 values of the slow component together with the variance of the fast component from a window around the same timepoint, with window size the same as the timescale. (Note the log-log scale of the axes, so that for a dependence of type *y* = *kx^n^*, the apparent offset in the log-log plot is defined by slope of the dependence *k*, and the apparent slope in the plot by the exponent *n*.) In agreement with our assumptions, the variances and the means are linearly dependent (*n* = 1), and in the case of raw photon counts, the dependence is exactly 1 (solid diagonal line *σ*^2^ = *μ*). In the standard mode with non-unit gain, the dependence is still linear, yet with a slope larger than one (dashed orange line). Points above the bulk in this view are due to windows with larger variance, and presumably connected with activity (at the appropriate timescale). Points below the bulk are due to undervariate windows. They are concentrated at high values and are due to saturated pixels. In the lower right plot we emphasise this fact by showing the points from saturated ROIs in red.

**Figure S6-4.**
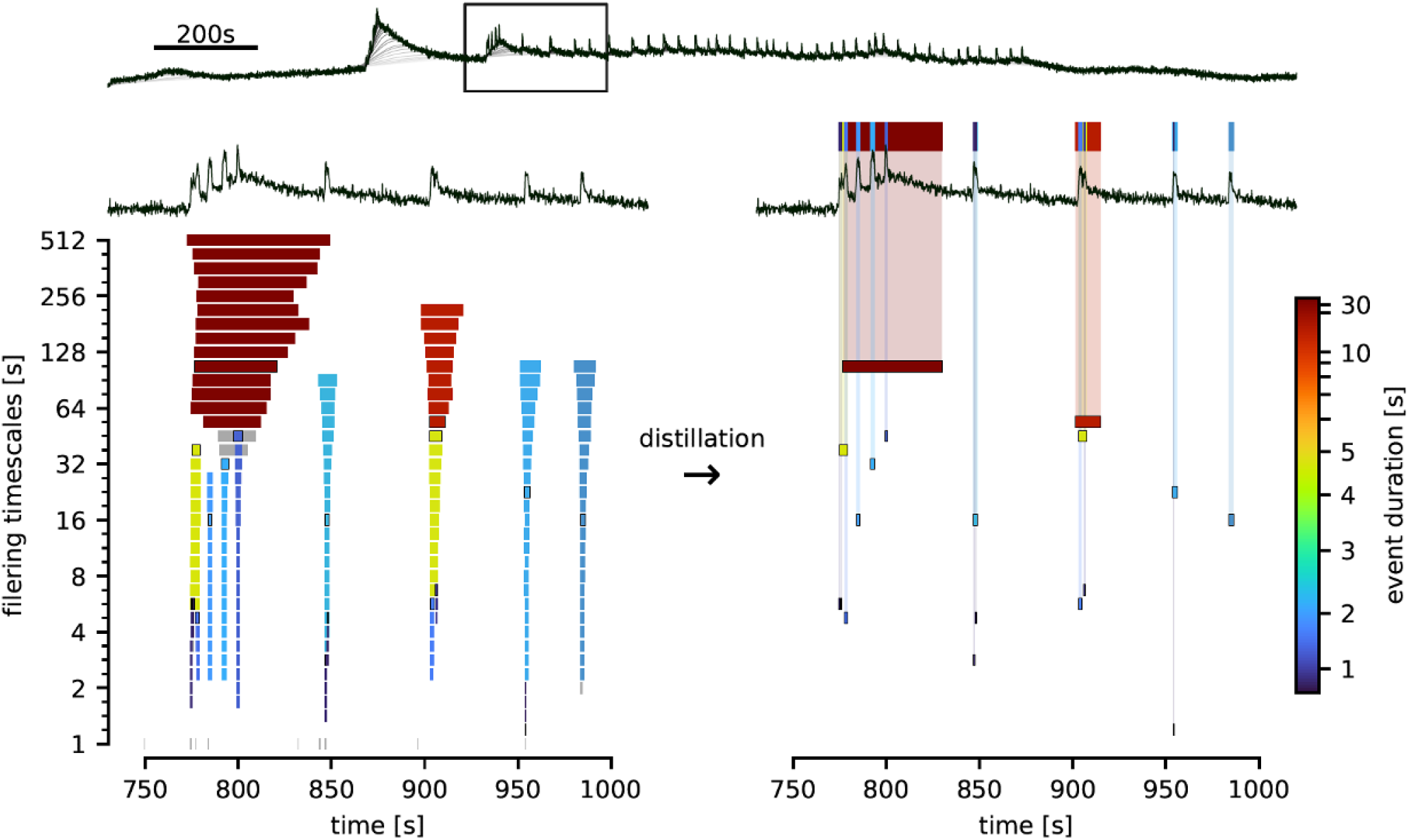
**A**, Example of a trace (black) and slow components in sequential filtering (shades of grey). **B**, Close-up view of the indicated small region from (A) and a representation of all candidate events in that region detected through sequential filtering. Each bar represents an event. Vertical position corresponds to the timescale at which the event was detected, the horizontal boundaries correspond to beginning and end time. In grey we show events that are deemed false positives, because they are detected at too few filtering timescales. Others events belong to groups of similar events which are then considered real; we show a boundary around the bar of the event closest to the one finally distilled. It’s time boundaries are set to median of the whole group, and the colour indicates its halfwidth. **C**, Distillation greatly removes redundancies and false positive candidates. Only robust events are selected for further analysis (see text for details). In the top part we represent the distilled events at single height, in a more compact view. We choose a colour code (mapping to a hill curve) which is nearly linear up to a few seconds, to emphasise the colour difference at the timescales most interesting for our analysis.

---

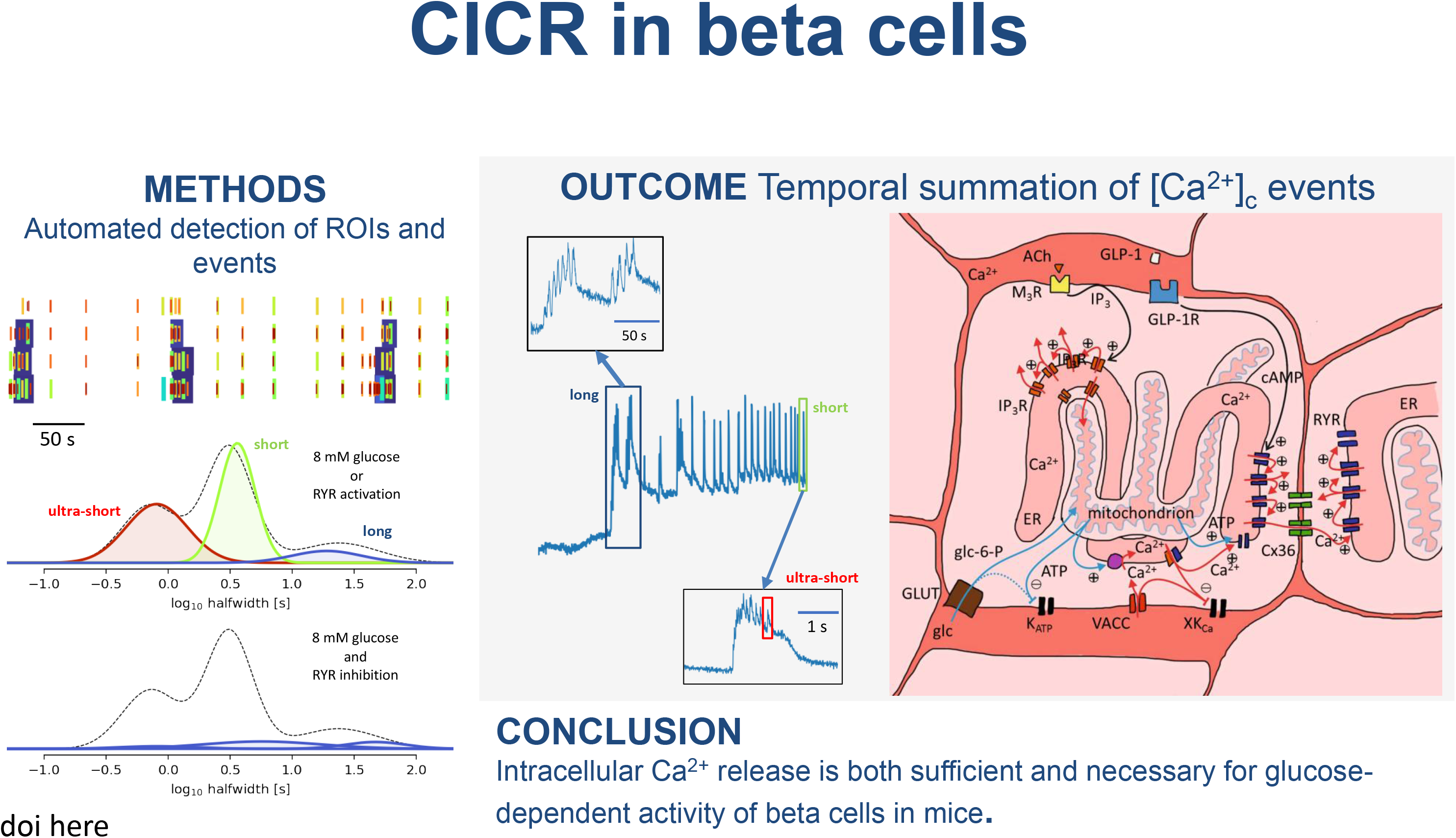

